# Deciphering functional dark matter: Machine and deep learning-based processing of protein embeddings enables targeted function discoveries

**DOI:** 10.64898/2026.05.29.728830

**Authors:** Sandra Wiegand, Anne-Kristin Kaster

**Affiliations:** Institute for Biological Interfaces 5, Karlsruhe Institute of Technology (KIT), 76344 Eggenstein-Leopoldshafen, Germany; Institute for Applied Biosciences, Karlsruhe Institute of Technology (KIT), 76131 Karlsruhe, Germany

**Keywords:** [NiFe] hydrogenases, [FeFe] hydrogenases, protein annotation transfer, alignment-free methods, sequence homology detection

## Abstract

The ever-expanding catalogue of uncharacterized proteins – the so called functional dark matter - poses a major challenge for biotechnological and biomedical exploitation. Functional assessment of most proteins is hindered by the technical limitations of annotation transfer and by the propagation of erroneous annotations in databases. The common denominator here is the reliance on sequence similarities. However, these become inaccurate below certain thresholds and can diverge even at sequence identities around 70%. To approach this challenge, we implemented a strategy using embeddings generated by protein language models for targeted function discovery (PE-TFD). Datasets of proteins representing target as well as non-target functions were used to train supervised learning models. The resulting ensemble models yielded interpretable prediction scores, enabling the exploration of databases without relying on multiple sequence alignments or structural information. We here tested PE-TFD for the discovery of novel hydrogenases as proof-of-concept, resulting in the novel discovery of 773 [NiFe] and 1,929 [FeFe] hydrogenases that were not detected by established sequence- or profile-based approaches. Structural analyses supported their non-random nature and further revealed a significant number of enzymes lacking prior functional annotation. Our framework therefore enables interpretable function discovery in large-scale datasets and the exploitation of functional dark matter.

**Graphical abstract:** 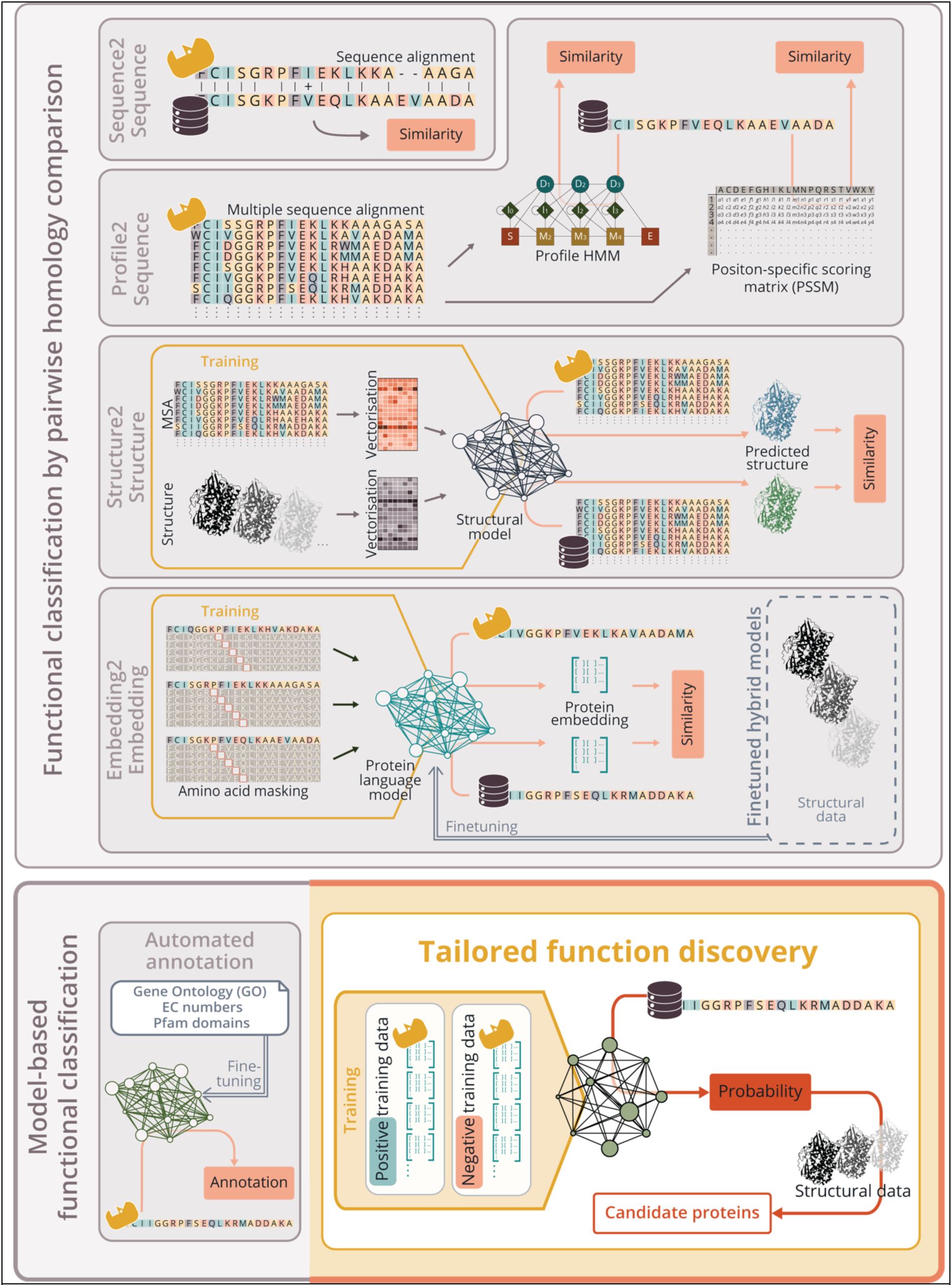

## Introduction

Proteins with unknown function are summarised by the term ‘functional dark matter’ (1). Their investigation is a crucial factor for *e.g.* understanding evolution, and exploiting microbial diversity for biotechnological advancements (2). While expanding sequence databases offer untapped possibilities, analysing and annotating the vast array of proteins remains challenging.

The most prevalent and fundamental method for identifying function is to address shared evolutionary origins by determining sequence similarities between characterized and uncharacterized proteins. Alignment algorithms enable searching public databases with customized queries. However, these are not able to reliably identify homologs with sequence identities below 30%, sometimes referred to as the *twilight zone*. (3, 4). More sophisticated profile-based methods, such as PSI-BLAST, which utilizes position-specific scoring matrices (PSSMs) (5) or profile hidden-Markov models (HMM), allow the detection of more remote homologs and have constituted the gold standard for homology searches for many years (6, 7). Both algorithms derive patterns from multiple sequence alignments (MSAs) of the target sequences and the position-specificity of HMMs, in particular, can lead to reliable predictions of homologous relationships as both the position within the sequence and the nature of adjacent regions are considered. Researchers can either generate custom profiles to search databases or utilize established resources such as Pfam, NCBIfam or others to characterise proteins or their domains (8–11). Profile-to-sequence methods are more sensitive in positive identifications within the *twilight zone* but are far from delivering highly reliable results (12, 13). Below 10% sequence similarity the so-called *midnight zone* begins (3), where even profile-based methods struggle to distinguish distant evolutionary relationships from random noise. The primary challenge is therefore not the absence of true positives, but the abundance of false positives.

The tertiary structure of functionally related proteins exhibits greater conservation than their amino acid sequences (14), therefore rendering it more suitable for functional homology detection in the midnight zone. Structure-based annotation approaches have existed for some time, but typically rely on the 3D crystal structures of protein domains (15, 16). However, only approximately 180.000 structures have been solved experimentally over the past 50 years, from which only about 12.000 proteins are unique based on their sequence identity. Luckily, structure-based homology searches for complete proteins have recently become viable (17, 18) and advances in deep neural networks have enabled protein tertiary structure modelling based on MSAs and structural information with unprecedented precision, generating over 200 million predicted 3D structures for analysis (19). Concurrently, newly developed algorithms for efficient structural alignment have made large-scale structure comparisons computationally feasible (20).

A significant challenge when analysing functional dark matter is the requirement for reliable MSAs, which are only achievable when sufficient sequences and structures with appropriate similarity ranges are available. As long as there are still novel amino acid sequences found through *e.g.* metagenomics and single cell data sets, there will always be a deficit of these required similar sequences. However, protein language models (pLM) now provide a tool to make predictions based on single sequences, thereby enabling alignment-free identification of remote homology. These models utilize the same transformer architecture employed in natural language processing and are trained to accurately predict amino acids based exclusively on sequence context (21–23). The internal sequence representations that are formed, so-called protein embeddings, are now able to capture evolutionary relationships (24). While embeddings provide a representation of tertiary structure, they alone are insufficient to fully determine the 3D conformation as they are missing information such as angles and distances. Therefore, pLMs have been fine-tuned to incorporate physical protein information and predict 3D protein structures (24, 25). Lately, several tools that infer remote homology between protein embeddings have become available, operating *via* nearest neighbour search (26), alignment of positional embeddings (27), or by providing input for structure or HMM-based comparisons (28). These approaches have demonstrated the capacity to identify distant homologs within the midnight zone of <10% sequence similarity (25). An interesting approach for direct annotation transfer the fine-tuning of pLMs or structural models with functional information. Numerous developments have emerged, enabling incorporation of GO terms, EC numbers, or data from CATH and Pfam (29–35). Some projects have even specialized on microbial annotation, focusing on GO terms relevant for prokaryotes, identifying antibiotic resistance genes and viral factors, as well as carbohydrate-active enzymes, and incorporating synteny information (36–39).

However, next to the required computational workload these approaches can suffer from transferring only positive data, resulting in high false positive rates (40). Here, we present a discriminative approach for customisable pLM representation-based exploration of protein databases. In brief, machine-learning (ML) or deep-learning (DL) based classifiers are trained on the embeddings of query proteins, which are divided into positive and negative input data. The generated model can then be used to identify proteins related to the positive data set in the queried protein databases. In contrast to previous studies, this method enables the tailoring of targeted searches for enzymes relevant to environmental microbiology, biotechnology and biomedicine. By not simply collecting all remote functional or structural homologues of a query protein but by incorporating prior knowledge to restrict and guide the output of the algorithm, a comprehensive search for previously unidentified biotechnologically exploitable enzymes is enabled.

We tested our framework on biotechnologically exploitable hydrogenases. Hydrogenases constitute a diverse group of metalloenzymes that are classified based on the metals found in their active sites: nickel-iron [NiFe] hydrogenases, iron-iron [FeFe] hydrogenases, and iron-only [Fe] hydrogenases. However, as the later differ from the other hydrogenases in several aspects and are specific to archaeal methanogenesis, they were excluded from the analysis. [NiFe] hydrogenases and [FeFe] hydrogenases catalyse the bidirectional conversion between molecular hydrogen and protons/electrons (41), a reaction that potentially enables the substitution of platinum-group metal-based catalysts in fuel cells (42, 43). After substantial research efforts in recent years, it has become evident that more stable hydrogenases are necessary to improve this process (44). The lack in identification of sufficiently stable candidates makes these enzymes therefore particularly well suited for testing novel discovery strategies (45). To identify promising, yet unannotated candidates, querying public databases for remote homologs of this protein family is inevitable. The application of the here developed PE-TFD (Protein Embedding-based Targeted Function Discovery) method allowed the successful identification of 773 [NiFe] hydrogenases and 1929 [FeFe] hydrogenases, which could not identified by the classical gold standard methods, thereby underscoring the potential of modern AI applications, and especially protein embedding-based approaches, to explore and illuminate functional dark matter.

## Methods

### Targeted function prediction

The study consisted of three main steps: (i) data selection and preparation, (ii) the training of diverse machine-learning models as well as a feed-forward neural network (FNN), and (iii) database screening using the trained models to predict novel candidates of the target proteins – in this case different classes of hydrogenases. The entire pipeline is implemented in Python, leveraging the PyTorch and scikit-learn libraries. All experiments were performed on a GPU server.

#### Data preparation

##### Protein embeddings

The Ankh_large model of the Ankh Transformer was used for generating protein embeddings (22). Sequences were processed in batches containing 100,000 to 350,000 tokens (including padding tokens), depending on sequence length. Sequences containing non-standard amino acids, pyrrolysine and selenocysteine, were excluded. The model produced embeddings with 1536 dimensions in latent space. To comply with the FNN design requiring fixed-size inputs, the mean representation of the last hidden state across the sequence was calculated for each protein. These representations were stored as gzipped JSON files and used as numerical input features for supervised learning.

##### Assembly of training data

For a balanced training, the precise separation of sequences into positive and negative datasets is crucial, as it can strongly influence the results. The positive dataset was derived from HydDB (46), a database exclusive to hydrogenase sequences. In addition, all sequences were curated by verifying their detectability using several major pre-computed HMM models for hydrogenase classes (8–10). The predictions were performed with hmmsearch (HMMER 3.1b2) (6), applying domain score–based cut-offs. For [NiFe] hydrogenases, all hits with the Panther database model PTHR42958 > 350 and/or the Pfam database model PF00374 > 250 were retained (9, 10). For the NiFe] hydrogenases, only the large subunits were used for the analysis, as they contain the active centre. For [FeFe] hydrogenases, sequences with the TIGRFAM database models TIGR04105.1 > 400 and/or TIGR02512.1 > 400 were kept (8). For [Fe] hydrogenases, which were only used as part of the negative dataset, all hits with TIGR01723.1 > 400 were included.

The Shannon index for the positive datasets was calculated upon clustering with MMseqs2 (v15.6f452) (47) with different minimal sequence identities per cluster.

The negative dataset was assembled from different sets of proteins. The set comprised random proteins from UniRef50 release 2024_01 (48). In addition, the set contained sequences annotated with EC numbers starting with EC 1.-.-.- (oxidoreductases), and had sequences with EC 1.12.-.- (acting on hydrogen as donors) annotations. Proteins annotated with EC 1.12.99.6 (hydrogenase (acceptor)) were explicitly excluded from the negative data. Furthermore, the datasets serving as the positive sets for the other two hydrogenase groups were also added as negative training data. All proteins with a size between 40 and 1000 amino acids were embedded as described above (*Protein embeddings*).

##### Explored databases

UniRef50 release 2024_01 (48) was downloaded, and all proteins with lengths between 40 and 800 amino acids were embedded as described above (*Protein embeddings*). To account for the possibility that proteins from the UniRef50 database were part of the training data, all protein instances present in the training dataset were systematically removed from the target database.

#### Modelling setup and training

##### Clustering and splitting of training and validation data

Given the limited availability of positive samples, a modified version of k-fold cross-validation was applied. Firstly, positive and negative datasets were clustered using the bisecting K-means algorithm to ensure balanced distribution in the folds (49). After clustering, the dimensionality of the data was reduced for Principal Component Analysis (PCA) and Uniform Manifold Approximation and Projection (UMAP) to allow for visualisation. The datasets were then partitioned into *k* folds, respecting the clustering structure. *k* models were trained per run, with each fold serving once as input for validation. Finally, all trained models were combined into an ensemble model that obtains final predictions by averaging outputs from the *k* models.

##### Feed-forward neural network (Deep-learning classifier)

The FNN was designed as a binary classifier with a two-dimensional output layer, followed by the calculation of probability values. The classifier had an input layer matching the dimensionality of the protein embeddings, followed by *n* fully connected hidden layers. Each hidden layer used a rectified linear activation function (ReLU) and dropout regularization. For optimization, the pipeline supports Stochastic Gradient Descent (SGD), Adam (50) and RMSprop.

To identify optimized model configurations, hyperparameters were tuned using the Optuna framework (51). Each trial sampled values for number of hidden layers (1 - 3), hidden layer widths (32 - 128), dropout rate (0.1 - 0.5), learning rate (10^−4^ - 10^−1^), optimizer type (Adam, SGD, or RMSprop), batch size (16 - 64), and number of training epochs (10 - 50). In addition, the Optuna trial cycle was repeated with tests for the effects of L1 (Lasso) and L2 (Ridge) regularisation, by optimising also for the L1 regularization coefficient (10^−6^ - 10^−2^) or the L2 weight decay (10^−6^ - 10^−2^), respectively. During each trial, the model was trained independently on all *k* folds and evaluated on the corresponding holdout fold. The optimization objective was the mean classification accuracy across all folds. During training, early stopping was applied based on validation loss with a patience criterion of 5 epochs. Finally, the probability values were determined by a softmax activation function.

##### Machine learning classifiers

Five supervised machine learning classifiers were used for the prediction of proteins from the learned embeddings: *k*-Nearest Neighbours (KNN), Gaussian Naive Bayes (NB), Support Vector Machines (SVM), Linear Regression (LR) and Random Forest (RF). Classifier hyperparameters were also trained using the Optuna framework (51). For KNN, the hyperparameter n_neighbors (1 - 20) was optimised across 50 trials. For NB, the hyperparameter var_smoothing (10^−9^ - 10^−1^) was optimised across 50 trials. For SVM, the hyperparameters for regularization C (10^−5^ – 10^2^) and kernel type (linear, rbf, poly) were optimised across 50 trials. For LR, the hyperparameters for regularization C (10^−5^ – 10^2^) and solver type (liblinear, saga, lbfgs) were optimised across 50 trials. For RF, the hyperparameters for the number of trees (n_estimators, 2 – 200) and the minimum number of samples required for splitting (min_samples_split, 2 – 20) were optimised across 50 trials.

### Benchmarking against other methods

#### Sequence-based searches

All amino acid sequences classified in HydDB (46) were used to perform DIAMOND (v0.9.14.115) blastp searches (52) against the target database in sensitive mode. The resulting hits were deduplicated, retaining only the hit with the highest bitscore for further analysis.

#### Profile-based searches

##### HydDB-based profiles

For each of the 38 HydDB classes, as well as for the three major hydrogenase groups, multiple sequence alignments were generated using Clustal Omega v1.2.4 (53). These alignments were used to build HMM profiles with hmmbuild (HMMER 3.1b2) (6). Additionally, PSSMs were generated using PSI-BLAST (BLAST 2.5.0+) (5). All generated models were used for prediction on the target database. As with the sequence-based searches, results were deduplicated, and the hit with the highest bitscore or highest domain score was selected for further analysis.

##### Public HMM-based profiles

For each of the two analysed hydrogenase groups, [NiFe] hydrogenases and [FeFe] hydrogenases, a defining set of domain and protein family models was manually compiled and fine-tuned on HydDB data (46). A major focus was to not only identify candidates but also to filter out proteins found to produce false positives in the context of hydrogenase detection, such as NuoD and Nar1p/Narf. These proteins are either components of respiratory complex I or are involved in iron-sulfur (FeS) cluster maturation, and they share a common evolutionary ancestor with NiFe hydrogenases and FeFe hydrogenases, respectively (54, 55).

[FeFe] hydrogenases were defined by hits against NF040762.1, NF040763.1, TIGR02512.1, TIGR04105.1, PF02256.21, or PF02906.18, but excluding those with hits to PTHR11615.SF124 or PTHR11615.SF258. [NiFe] hydrogenases were characterized by hits against NF033181.1, NF040828.1, NF040832.1, NF041785.2, TIGR03295.1, PTHR42958, or PF00374.23, while excluding hits to PF00329.23, PF00346.23, PTHR11993, or TIGR01962.1. The corresponding data were obtained from Pfam v36.0 (9), PANTHER 19.0 (10), and the NCBI Protein Family Models database NCBIfam (8). Where necessary, HMMs were built using hmmbuild (HMMER 3.1b2) (6). The prediction was performed with hmmsearch (HMMER 3.1b2) (6).

#### Structure-based searches using Foldseek

For structure-based searches, pre-folded protein structures corresponding to HydDB-listed proteins were retrieved from AlphaFoldDB (19) in June 2024. To reduce the number of search queries, proteins were clustered within their respective groups using MMseqs2 v15.6f452 linclust (56) with identity thresholds of 0.5 for [NiFe] and [FeFe]. The representative structures (Table S1) were used to query the pre-folded AlphaFold/UniProt50 v4 database using the Foldseek webserver in TM-align mode (20). The results were deduplicated based on the highest score and cross-referenced to UniRef50 v2024_01 using BLASTP (BLAST 2.5.0+) (57) to enable comparison with the other methods.

### Evaluation

#### Classification

Hits from the public HMM-based profiles were classified with the cut-off defined in Table S2. For [NiFe] hydrogenases, hits obtained from sequence-based and HydDB profile-based (best domain) pairwise comparisons were classified as *soft* (bitscore >50), *good* (>100), or *excellent* (>300) candidates. For [FeFe] hydrogenases, hits from sequence-based comparisons were classified as *soft* (bitscore >50), *good* (>300), or *excellent* (>350), whereas HydDB profile-based (best domain) pairwise comparisons used thresholds of >50 for *soft*, >175 for *good*, and >300 for *excellent* candidates. For Foldseek profile-based results, hits with a bitscore >30 were considered soft candidates, >65 as good candidates, and >90 as excellent candidates. Thresholds for each ML/DL classifier were determined individually for each group and model. For all methods, false positives were identified using hmmsearch with public HMM profiles as described in Public HMM-based profiles: ([NiFe]) hits against PF00346, PTHR11993, or TIGR01962 with scores >250, ([FeFe]) hits against PTHR11615.SF124 or PTHR11615.SF258 with scores >350. Positive, unique results were cross-checked using InterProScan v5.75-106 (58).

#### Structure comparisons

The structures of positive, unique results were predicted using ESMFold (23). These predicted structures were compared with reference structures using Foldseek v9.427df8a (20). The reference structures included: (i) HydDB entries with experimentally determined structures, (ii) HydDB entries with AlphaFoldDB (19) predictions and (iii) HydDB entries folded for this analysis with ESMFold.

## Results and discussion

To fully exploit the potential of microbial diversity harboured in functional dark matter, it is essential to establish techniques that enable improved transfer of functional knowledge. Automated annotation transfer in prokaryotes remains an ongoing challenge, primarily due to the limitations of existing annotation systems. Gene Ontology (GO) terms, for instance, were originally developed for functional annotation in eukaryotes (59), and bacteria and archaea are still largely underrepresented (60, 61). Although attempts have been made to incorporate prokaryotic information, recent efforts have been emphasising on pathogenicity (62). Annotation methods for prokaryotes typically rely on transferring functional information from well-characterised proteins to homologs with a sufficient degree of similarity. However, annotation transfer suffers below sequence identities of 70% and becomes unreliable once sequence similarities drop below 30% (3, 63).

To address these limitations, we implemented a function discovery strategy directly applicable to prokaryotic proteins that leveraged protein language model (pLM) embeddings as the central representation of protein features. pLM embeddings have been shown to outperform annotation transfer methods based on raw sequence similarity or MSA-based methods, particularly for proteins with very low (<20%) sequence identity (26, 64). This is especially relevant when exploring the functional dark matter of prokaryotic genomes, where evolutionary distances are large and high-quality MSAs are difficult or impossible to obtain (65). While structure-based methods are more accurate, they are often limited by the availability of structural data, which are frequently missing and costly to obtain for many prokaryotic proteins (66). Even single-sequence approaches such as ESMFold offer only limited relief, as they are ultimately trained on existing structural data and therefore perform best for proteins with close structural homologs (66, 67). Given these limitations, pLM embeddings offer a robust alternative that helps mitigate overfitting (68). In the context of this work, this was particularly advantageous, as our objective was not to assign highly specific annotations but to compile a broad list of candidate proteins that fall in a defined functional space.

Another limiting aspect that was addressed was the one-way comparison of uncharacterized proteins to known representatives of established protein families as mean of function transfer. When such analyses are performed within the *midnight* and *twilight* zones of sequence similarity, the major cause of incorrect predictions is not the loss of true hits through false negatives, but the abundance of false positives (3). To overcome this problem and also use the possibility to precisely define the functional space of interest, we trained supervised machine-learning and deep-learning classifiers using both positive and negative datasets. Five classical ML algorithms were employed alongside a DL feed-forward neural network.

### Compilation of the training data sets

Here, we present a protein embedding-based targeted function discovery (PE-TFD) approach to explore public databases for the presence of yet unidentified [NiFe] and [FeFe] hydrogenases as as a proof of concept. The underlying idea was to identify hydrogenases that may have escaped detection by classical sequence- and profile-based annotation methods due to low sequence identities, despite the fact that homologs can exist without detectable sequence similarity (69, 70). The identification of proteins that nonetheless appear to be similar to known hydrogenases in embedding space would expand the pool of candidate hydrogenases and may reveal enzymes with novel functional properties. One such feature of interest is *e.g.* high O₂ tolerance, a prerequisite for hydrogenases intended for use in fuel cells (71, 72). While pronounced O₂ tolerance is mainly associated with [NiFe] hydrogenases, [FeFe] hydrogenases have on the other hand been reported to exhibit higher turnover frequencies (73). As these two hydrogenase classes are not phylogenetically related and do not share detectable sequence similarity (54), both groups were included in our analysis to avoid overlooking potential functional novelties. [Fe] hydrogenases were not included in the analysis, as they do not contain nickel or iron-sulfur (FeS) clusters (74) and require methenyltetrahydromethanopterin for H_2_ activation. In contrast, [FeFe] and [NiFe] hydrogenases actively catalyse the H_2_/H_2_O exchange in the absence of exogenous electron acceptors (75).

Based on the laid out considerations, a dedicated prediction framework was established. To this end, the PE-TFD classifier models were trained in an initial step. Model training included an evaluation phase to assess the ability of the classifiers to generalize from the training data to previously unseen proteins. This required the assembly of a reliable and well-curated training dataset. Owing to their lack of evolutionary relatedness, [NiFe] and [FeFe] hydrogenases were treated as separate classification tasks and trained independently.

HydDB is a database containing 3248 annotated catalytic subunits categorized into 29 [NiFe] and 8 [FeFe] classes (46). These classes were populated using a sequence similarity network constructed from alignment distances (46). For model training, the [NiFe] and [FeFe] datasets were refined by mapping the respective signature HMMs required to define each main class. For [NiFe] hydrogenases, PTHR42958 and PF00374 were used, whereas TIGR04105.1 and TIGR02512.1 were applied for [FeFe] hydrogenases. In total, the positive training sets comprised 938 [NiFe] and 849 [FeFe] protein entries (Table S3). After clustering at 70% sequence identity, the datasets formed 222 and 391 clusters, respectively. The corresponding Shannon evenness values of 0.86 and 0.94 indicate a balanced representation of protein sequences within each dataset (Table S4).

Negative training sets were assembled from randomly selected protein sequences as well as proteins annotated as oxidoreductases (EC 1.-.-.-) and oxidoreductases acting on hydrogen as a donor (EC 1.12.-.-). In addition, each negative set included the positive training sequences of the respective other hydrogenase group as well as [Fe] hydrogenases. This strategy ensured that the classifiers learned to distinguish between the different hydrogenase classes. Overall, each group’s training set contained approximately fivefold more negative than positive sequences, providing sufficient class balance for robust model training (Table S3).

### Validation and testing setup

As outlined above, it is essential that the constructed models are rigorously evaluated. Usually, in machine learning workflows a dedicated test dataset is withheld during training and used only to evaluate the generalisation capability of the model - that is, its ability to identify suitable candidates in previously unseen data. It is crucial that this test set does not contain proteins overly similar to those in the training set and that the two sets are strictly kept separate; otherwise, the evaluation would primarily reflect the model’s ability to recognize near-duplicates rather than its true predictive capacity (70).

To prevent the described data leakage but also maximise the use of the scarce biological data, we employed *k*-fold cross-validation with ensemble averaging (Figure 1) (76–78). In a first step, the embedded data were clustered using bisecting *k*-means to group similar sequences together. With these clusters, *k* folds were generated. Then, *k* separate models were trained, with each fold serving as the holdout test set exactly once, while the other folds were used for training. This procedure resulted in *k* independently trained models; their prediction probabilities were averaged to produce an ensemble output. This strategy ensures that closely related proteins are not spread across training and test sets and maximizes the use of every data point as each protein contributes to both training and testing.

**Figure 1.**
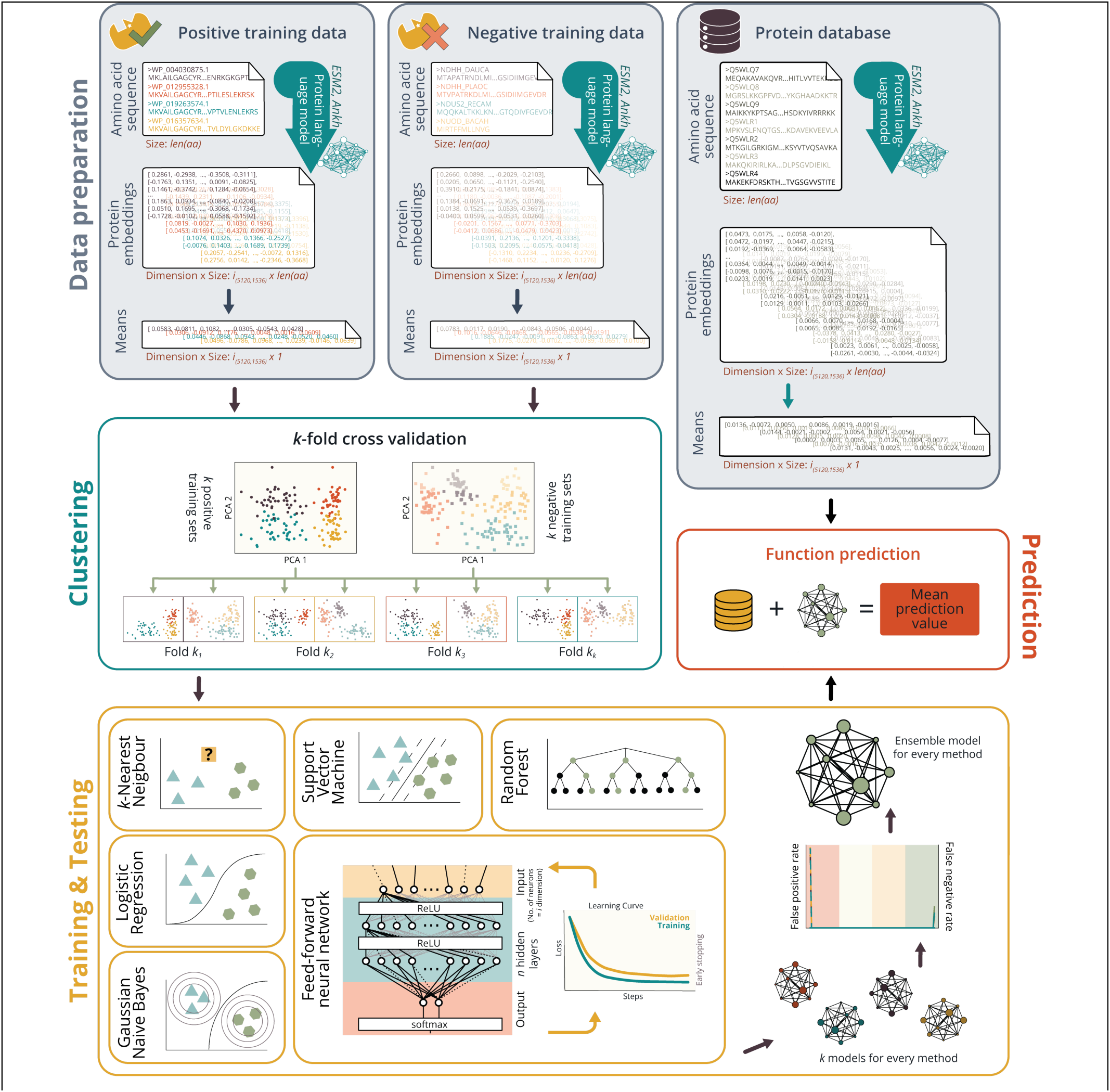
Protein embedding-based targeted function discovery (PE-TFD) workflow. Positive and negative training datasets were selected, embedded using a protein language model, and reduced by averaging the final hidden layer. Embedded proteins were then clustered and partitioned into *k* folds. During training, each of the employed machine-learning (Logistic Regression (LR), Gaussian Naive Bayes (NB), *k*-Nearest Neighbours (KNN), Random Forest (RF), Support Vector Machines (SVM)) and deep-learning classifiers (Feed-forward neural network (FNN)) were trained independently on each of the *k* folds and evaluated on the corresponding hold-out set. Upon inspection, the models passing the performance test were combined into an ensemble model. For the prediction, these ensemble models were applied to an embedded target database, and mean prediction value for each entry in the database were computed.

For training the neural network, an additional validation step was required. After training but prior to testing, each fold’s training subset was further split, reserving 15% as a validation set. This validation set was used exclusively for hyperparameter tuning and early stopping (*see below*). Testing was then carried out on the designated test folds, analogous to the machine-learning models.

### Modelling

The supervised learning models constructed in this study (Figure 1) were designed as binary classifiers to distinguish between negative and positive instances within each analysed dataset. To evaluate the modelling strategies for the targeted prediction of enzymes with hydrogenase activity, several algorithms were implemented to enable a systematic comparison of performance.

The machine-learning (ML) models included Logistic Regression (LR), Gaussian Naive Bayes (NB), *k*-Nearest Neighbours (KNN), Random Forest (RF), and Support Vector Machines (SVM). As a deep-learning (DL) method, a feed-forward neural network (FNN) was trained. These models differ in learning behaviour as well as interpretability, and achievable accuracy (78). According to the No-Free-Lunch theorem, no single algorithm is universally superior and performs optimally across all possible datasets (79). It was therefore essential to evaluate several modelling strategies.

LR was chosen for its ability to perform well on smaller datasets and because it provides interpretable probability outputs by design (80). NB assumes conditional independence of features within each class; nonetheless, it often performs robustly even when this assumption is violated (81). KNN is distinct in that it requires no conventional training phase. The training data constitute the model itself, and predictions are based on neighbourhood structure (79). SVMs construct a separating hyperplane between classes. They are computationally efficient but require the additional production of probability estimates. RF models learn the importance of individual features and permit interpretable, human-readable inspection of decision trees (70).

Neural networks, while more complex, can produce more accurate predictions when sufficiently trained. What distinguishes ML from DL is not the use of a neural network *per se* but the presence of multiple hidden layers (78). Although “deep learning” does not have a strict layer-count definition, here we use the term for networks with 2 or more hidden layers. The feed-forward neural network used in this study consisted of an input layer corresponding to the 1,536-dimensional pLM embeddings generated by the Ankh transformer (22), followed by *n* fully connected hidden layers with rectified linear unit (ReLU) activations and dropout regularisation. For optimisation Stochastic Gradient Descent (SGD), Adam, and RMSprop were tested. Additional hyperparameters subjected to tuning included the number and width of hidden layers, dropout rate, learning rate, batch size, and number of training epochs.

To prevent the model from overfitting, which is a modelling error that occurs when a model learns the training data too closely, multiple regularisation strategies were implemented. Dropout (82) was applied to both input and hidden layers, randomly disabling a fraction of neurons at each training step to prevent the model from focusing on a single neuron. Early stopping halted training when the validation loss failed to improve for five consecutive epochs. Both L1 and L2 regularisation of the loss function were examined to penalise overly complex models (83). Cross-entropy loss combined with softmax activation was used to compute stable class probabilities and gradients during optimisation. Learning curves (Figure 1) were monitored to ensure appropriate training behaviour, with ideal models showing rapid decreases in both training and validation loss. To identify suitable model configurations for all six modelling approaches, hyperparameters were automatically optimised.

### Training and Evaluation

Training for the [NiFe] and [FeFe] hydrogenase prediction models began with an initial focus on the evaluation of the behaviour of the training data. After clustering, the dimensionality of the data was reduced to visually inspect the adequate separation of clusters. A suitable number of clusters (*j)* was determined for both enzyme classes, and several training runs were performed using 0.5*j* to 2*j* clusters and *k* folds ranging between *j/3* and *j* to assess how the number and ratio of clusters and folds influenced model performance. Based on inspection of the resulting cross-validated models, the optimal and most robust results were gained with 10 clusters and 5 folds for the [NiFe] hydrogenase training and 12 clusters and 6 folds for the [FeFe] training. The optimised hyperparameters associated with these cluster configurations for all six algorithms are summarised in Table S5.

Model performance was primarily evaluated on probability threshold-dependent plots of the false positive rate (FPR) and false negative rates (FNR) ([NiFe] hydrogenases Figure 2a, [FeFe] hydrogenases Figure S1-S3). A false positive (FP) corresponds to a negative sample incorrectly predicted as positive, while a false negative (FN) represents a missed positive sample. The corresponding rates were calculated by dividing FP and FN by the total number of true negatives and true positives, respectively. These two values are the complements of specificity and sensitivity, which measure true detection rates rather than reflecting error rates.

**Figure 2.**
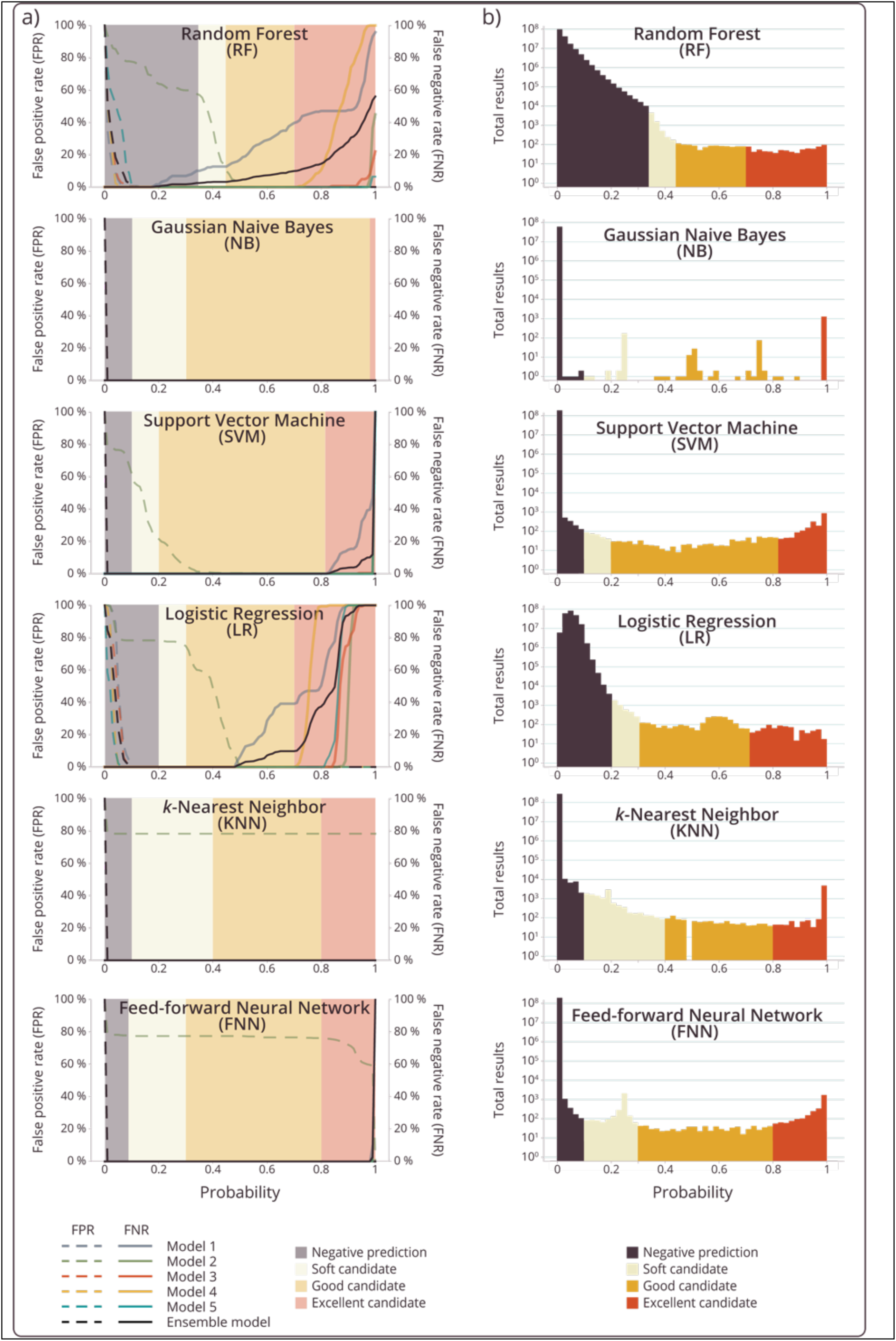
Performance of each classifier during testing and prediction - [NiFe] hydrogenases. **a)** False positive rates (FPR) and false negative rates (FNR) across probability thresholds. Each panel shows the FPR (dashed lines, left y axis) and FNR (solid lines, right y axis) for models 1 to 5 across probability thresholds for one of the applied classifiers. In addition, the rates of the ensemble model (models 1,3,4,5) are given. The curves inform the selection of probability thresholds that balance sensitivity and specificity for identifying novel [NiFe] hydrogenase candidates. **b)** Number of total prediction results across probability thresholds. Distribution of predicted probabilities for all proteins in UniRef50 generated by each PE-TFD classifier. The background shading for a) and b) indicates the probability ranges used to define prediction categories: negative prediction (red), *soft* candidates (light orange), *good* candidates (yellow), and *excellent* candidates (green).

For [NiFe] hydrogenases (Figure 2a), FPR values are high at low probability thresholds, as nearly all instances are classified as positive when the threshold approaches zero. As the threshold increases, FPR decreases for all algorithms with one notable exception: model 2 displays persistently elevated FPRs for all but the NB algorithms, predicting many false positives originating from the negative dataset.

This misclassification arises from the data composition and relates to the special case of the respiratory NADH-quinone oxidoreductase subunit NuoD, which is structurally similar to hydrogenase large subunits. [NiFe] hydrogenases contain a large catalytic subunit harbouring the active-site [NiFe] cofactor, which is the primary target of this analysis, and a small subunit containing one or more FeS clusters (55). Both the large and small hydrogenase subunits share homology with the NuoD and NuoB subunits of respiratory complex I, respectively (55). Earlier evolutionary models proposed that membrane-bound hydrogenases were the progenitors of complex I (69) but recent analyses suggest that complex I and modern [NiFe] hydrogenases share a common ancestor, likely an Mrp-type Na⁺/H⁺ antiporter (84).

The behaviour of model 2 can be attributed to the clustering of the dataset, as the holdout negative fold reserved for model 2 testing contained many, if not all, NuoD-like proteins. As a result, the model was not exposed to these closely related non-hydrogenase proteins during training, impairing its ability to distinguish them from true hydrogenases during testing. Despite this limitation, the SVM, LR and RF models were able to manage this challenge reasonably well at mid-range probability thresholds. In contrast, KNN and the neural network model were unable to recognize the relevant evolutionary relationships without the explicit presence of these sequences in the training data. To acknowledge the importance of these data during training and to avoid a consistently high FPR in the ensemble models, model 2 was excluded from all subsequent analyses, allowing us to pursue a bespoke solution for NiFe hydrogenase data handling (70).

For [NiFe] hydrogenases, the determined FNRs behave inversely: at low probability thresholds, FNRs remain low, as the model is permissive and labels most proteins as positive. As the probability cutoff becomes more stringent, FNRs increase gradually, reaching 100% for LR, RF, SVM and FNN. This increase enables the determination of thresholds for which the predictions are reliable but more conservative, thereby overlooking candidates (*excellent* candidates). This does not interfere with the goals of this study, as we primarily aim to identify as many novel candidate sequences as possible with high confidence; candidates missed at high thresholds will still be captured when applying lower probability cutoffs (*soft* and *good* candidates).

As the probability cutoff becomes more stringent, FNRs increase, reaching 100% for LR, RF, SVM, and FNN. This increase enables the identification of thresholds above which predictions are reliable but more conservative, thereby overlooking some candidates (i.e., excellent candidates). This behaviour does not interfere with the goals of this study, as our primary aim is to identify as many novel candidate sequences as possible with high confidence; candidates missed at high thresholds will still be captured when applying lower probability cutoffs (soft and good candidates). The boundaries of these classification ranges are visualised in Figure 2a by the background shading.

### Targeted prediction of hydrogenases

Upon finalisation of the six ensemble models, predictions for all six PE-TFD classifiers were carried out on all proteins in the UniRef50 database with lengths between 40 and 1000 aa, comprising a total of 59.9 million proteins. Each classifier produced a probability score for every embedded protein, as shown in Figure 2b. The different distributions of the prediction results reflect the distinct behaviours of the FNR and FPR curves, particularly for the NB and KNN classifiers, and illustrate that some classifiers are more rigid than others in shifting predictions towards the borders of the probability range. The introduced classification scheme of soft, good and excellent proteins captures these tendencies and renders the results comparable (Figure 3a), yielding between 1553 and 9014 positive predictions depending on the method. As intended by the architecture of the ensemble model, which only included models that had NuoD in the training data, only very few of the proteins were classified as NuoD by publicly available HMMs. However, a noticeably large proportion of NuoD proteins appeared within the soft candidate group of the FNN model, suggesting that this classifier has a broader detection range at lower thresholds and consequently yields soft candidate predictions with lower confidence.

**Figure 3.**
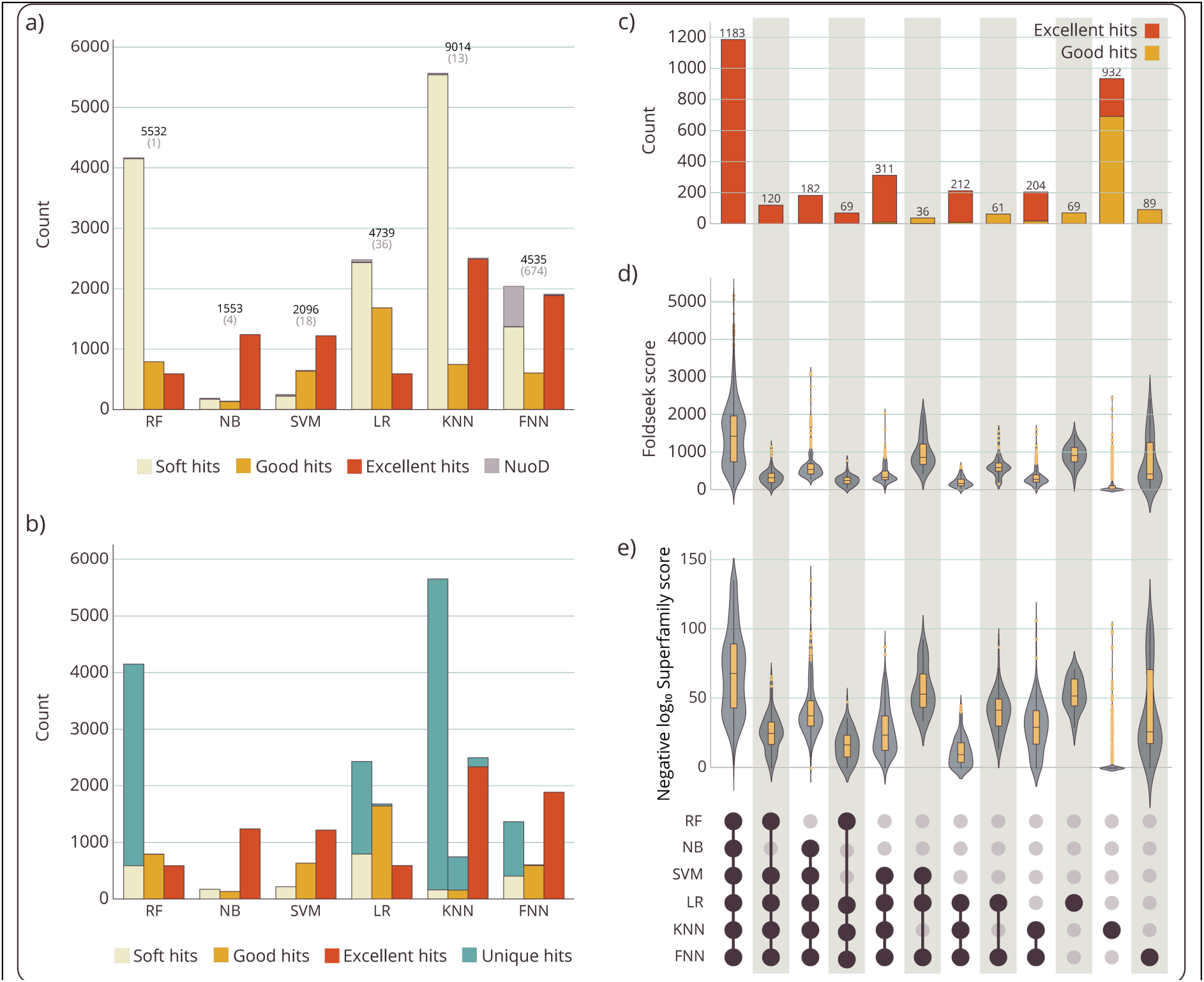
PE-TFD classifier performance on [NiFe] hydrogenase-aimed predictions. **a)** Numbers of *soft*, *good*, and *excellent* candidates predicted by each classifier with Fraction of incorrectly identified NuoD proteins. **b)** Proportion of candidates detected exclusively by each classifier. **c-e)** Intersection analyses of good and excellent candidates and the six applied ML/DL classifiers. Combinations with fewer than 10 proteins were omitted for clarity. **c)** Number of hits per combination. **d)** Foldseek probabilities of ESMFold-predicted structures of PE-TFD candidates and ESMFold-predicted structures of reference [NiFe] hydrogenases from HydDB for being homologous. **e)** Confidence scores of hydrogenase-associated structural domains of the Superfamily database in PE-TFD candidates.

Another important aspect of the predictions is whether a given candidate is supported by more than one classifier (Figure 3b). This was frequently not the case for soft candidates, especially those predicted by RF, LR, KNN, and FNN, for which the majority of these proteins were not recovered by any other classifier. This observation further highlights the lower reliability of the soft candidate category, and therefore this category was omitted from subsequent analyses.

Across the good and excellent categories, a total of 3517 protein hits were retained for further evaluation (Figure 3c). Of these, 66% were classified as excellent by at least one classifier. While a total of 34% of all hits were detected by all PE-TFD classifiers, 69% were identified by at least two classifiers, leaving the other 31% to have been identified by only a single classifier.

When comparing the individual classifier results, it becomes clear that NB and RF yield the lowest numbers of good and excellent hits, with 1375 and 1383 proteins, respectively. These are followed by SVM with 1857 hits, LR with 2274 hits, FNN with 2494 hits, and finally KNN, which produces the highest number at 3240 hits. This ranking is also reflected in the overlap across classifiers: hits produced by RF, NB, and even SVM are almost always supported by at least two additional classifiers, while LR, FNN, and KNN also generate subsets of hits that receive support from fewer other methods.

All 3517 hits were further examined for structural features by comparing their ESM-folded structures against ESM-folded structures from all [NiFe] hydrogenases available in HydDB (Figure 3d) and by analysing their structural domains using the Gene3D and SUPERFAMILY databases, which are based on the CATH and SCOP classification systems and employ HMM-based identification of known structural domains (Figure 3e + Figure S4). The strongest trust values were consistently assigned to proteins detected by all PE-TFD classifiers. High trust values were also observed for groups dominated by LR and FNN, including their overlap with SVM. In contrast, groups that include KNN-specific hits stand out due to markedly lower trust values - the group of hits only found by KNN even yields by far the lowest median and third quartile values. This suggests that the KNN model identifies a set of candidates that is distinct from those found by the other methods; however, whether these represent genuine hydrogenases or false positives remains unresolved. To eliminate the risk of bias from possible false positives, KNN-predicted candidates were not considered during benchmarking.

### Benchmarking of the PE-TFD predictions

The next stage in evaluating the targeted machine-learning classification (PE-TFD) of hydrogenases was benchmarking the predictions discussed above (Figures 2+3). These predictions were compared against functional homology assignments obtained using either established sequence- and profile-based approaches or 3D structure-based methods. Sequence-based similarity searches were performed using BLAST (*Seq2Seq*). Profile-based methods employed profile HMMs and PSSMs constructed from multiple sequence alignments of the respective HydDB classes (*HydDB-HMM2Seq* and *HydDB-PSSM2Seq*). In addition, publicly available HMM resources including Pfam, TIGRFAM, PANTHER, and NCBIfam were queried (*Public HMM2Seq*). For structure-based comparisons, pre-folded protein structures corresponding to HydDB-listed sequences were retrieved from AlphaFoldDB and used to search the pre-folded AlphaFold/UniProt50 database (*Struc2Struc*).

Across all approaches, the number of good and excellent predictions fell within a comparable range, spanning 1342 to 4106 hits. As an initial benchmark, the extent of NuoD misidentification was quantified for each method (Figure 4a). A substantial rate of incorrect NuoD assignment among good and excellent candidates was observed for sequence-, profile-, and structure-based methods, with error rates of 27%, 25%, and 38%, respectively. Misannotation was absent only in methods in which NuoD detection was explicitly incorporated - either through targeted training of the ML and DL models or through the usage of several public profile HMMs that already include NuoD as a searchable instance (*Public HMM2Seq*). At first glance, the results underscore a central limitation of traditional homology-based approaches: insufficient discrimination between true positives and closely related false positives. While the lower number of candidates identified by PE-TFD may of course reflect the omission of some true hydrogenases, the reported lack of discrimination between unconserved protein functions at high sequence identities (63) is also very likely to explain the substantially higher hit counts observed for sequence- and profile-based methods.

**Figure 4.**
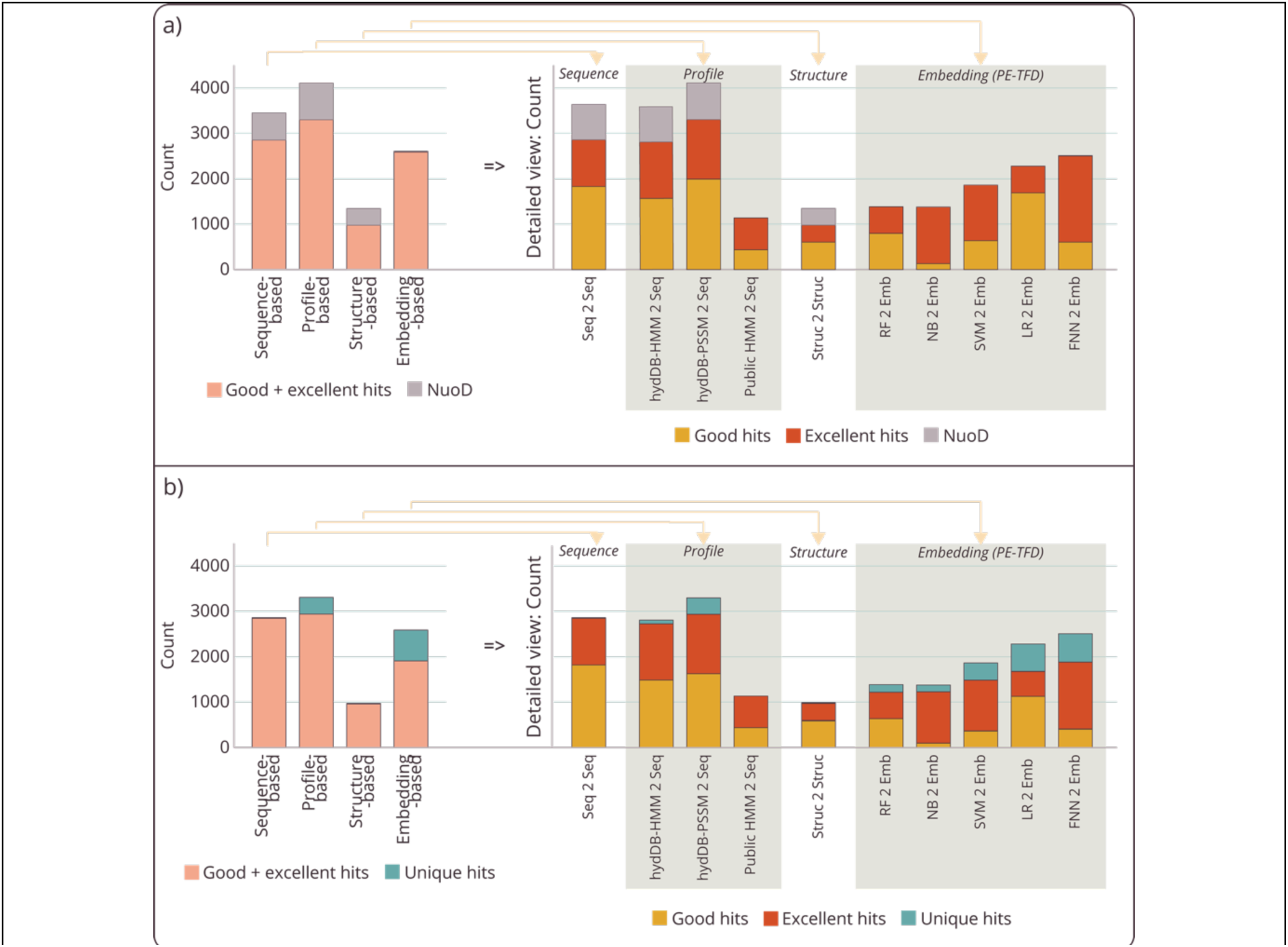
Benchmarking PE-TFD results against sequence-, profile-, and structure-based approaches. **a)** Fraction of incorrectly identified NuoD proteins among good and excellent candidates produced by different method categories, including sequence-based (*Seq2Seq*), profile-based (*HydDB-HMM2Seq*, *HydDB PSSM2Seq*, *Public HMM2Seq*), structure-based (*Struc2Struc*), and protein embedding-based (RF, NB, SVM, LR, FNN) approaches. **b)** Proportion of candidates detected exclusively by each method category. KNN predictions were not included in the analysis.

In a second step, the novelty of the detected candidates was assessed (Figure 4b). The three categories of benchmarking methods showed only limited capability to identify proteins not detected by any of the other methods. The fraction of such unique identifications ranged from 0.4% for *Seq2Seq* to 11% for the profile-based approaches. This observation is partly attributable to the smaller number of positive hits generated by *Struc2Struc* and *Public HMM2Seq* but is also derived from the extensive overlap between sequence- and profile-based predictions (Figure 5a). In contrast, ML- and DL-based predictions yielded a substantially higher fraction of unique hits not identified by the benchmarking tools. Unique candidates were particularly enriched in the outputs of the SVM, FNN, and LR classifiers, accounting for 20%, 25%, and 27% of their predictions, respectively. However, as shown in Figure 3c, overlaps indeed exist among the unique hits produced by the PE-TFD catalogue.

**Figure 5.**
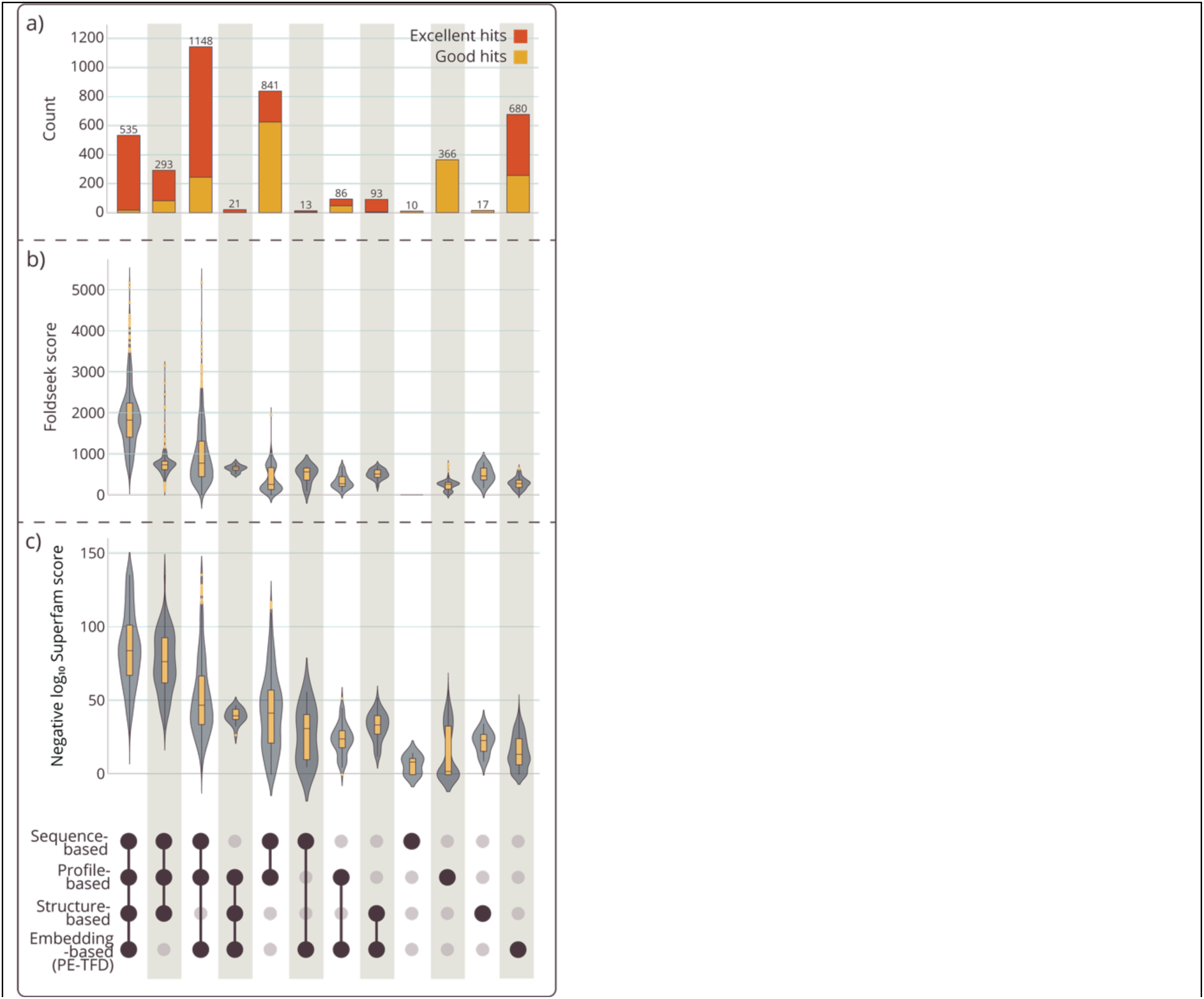
Intersections and structural breakdown of PE-TFD and benchmarking results. Combinations with fewer than 10 proteins were omitted for clarity. **a)** Number of hits per combination. **b)** Foldseek probabilities of ESMFold-predicted structures of PE-TFD candidates and benchmarks and ESMFold-predicted structures of reference [NiFe] hydrogenases from HydDB for being homologous. **c)** Confidence scores of hydrogenase-associated structural domains of the Superfamily database in PE-TFD.

From an overall perspective, only about half of the 4121 good and excellent candidates found in total were identified by at least three methods. Notably, the majority of these consensus predictions belonged to the excellent category. Structural comparison of these candidates with HydDB-listed hydrogenases revealed that high-confidence structural similarity scores (median score of 1825) were observed exclusively for proteins detected by all methods (Figure 5b). Candidates identified only by at least three methods showed median scores above 600. A consistent trend was observed for the presence of hydrogenase-associated structural domains (Figure 5c + Figure S5). For candidates unique to PE-TFD, both structural alignment scores and hydrogenase-associated domain scores were generally low, yet clearly above zero. This supports the non-random nature of the PE-TFD predictions, as *e.g.* the scores for the discarded KNN-derived candidates were drastically lower (Figures 3d+e).

One component of the benchmarking was to evaluate whether the structure-based, Foldseek-supported AlphaFoldDB search could serve as an alternative approach for AI-assisted prediction in functional dark matter. The results showed that the structure-based approach yielded fewer hits overall and, in particular, identified only a small number of candidates not detected by the two gold-standard methods (sequence- and profile-based searches), with only 110 such cases (Figure 4b + Figure S6). In contrast, PE-TFD identified 680 [NiFe] hydrogenase candidates not detected by any of the sequence- profile- or structure-based analyses. When restricting the comparison to the gold-standard methods of sequence and profile-based approaches, 773 proteins were classified as novel, accounting for almost 20% of all the 3517 proteins detected by these three methods (Figure S7). This clearly demonstrates the advantage of the protein embedding-based approach in uncovering additional candidate proteins. In a nutshell, the results for the [FeFe] hydrogenases were 1929 unique PE-TFD predictions, out of 3649 total embedding-based predictions (Figures S2+S3).

Finally, the nature of the 680 PE-TFD-unique [NiFe] hydrogenase candidates was examined. Obviously, they have no high sequence similarities detectable by sequence-o r profile-based methods as they were classified as *unique*. Above mentioned was their additional lack of high structural similarities to other hydrogenases as can be seen from the results of the structural similarity scores and the presence of hydrogenase-associated structural domains (Figure 5b+c). InterPro annotations exceeding the mentioned structural domain predictions were available for 41% of these proteins, and all such annotations were related to hydrogenases or respiratory complex I. However, in these cases confidence scores were consistently low, resulting in approximately half of these proteins being discarded and the remaining half classified as soft candidates. The examination of NCBI annotations revealed that 34% of the proteins were annotated as hydrogenases, often supported by corresponding InterPro results. A further 20% were annotated as maturation proteins, although InterPro provided no clear evidence supporting this assignment in all but one case. While a minority of proteins carried diverse alternative annotations, 37% were labelled as hypothetical proteins, underscoring the potential of modern AI methodologies, and especially protein embedding-based approaches, to explore and illuminate functional dark matter.

## Conclusion

Functional prediction of microbial proteins is inherently challenging, as similar functions do not necessarily go along with high sequence or structural similarity. Functions can be conserved despite sequence dissimilarities, and even small natural sequence variations observable within enzymatic families can have poorly understood effects on function (85). This knowledge gap is reflected in benchmarking studies of commonly used databases, which report misannotation rates for enzymatic protein families exceeding 80% (86). Moreover, it has been demonstrated that nearly 20% of the members of a well-defined enzyme class lack any detectable sequence or structural similarity to experimentally characterized representatives (87). In addition to errors introduced through information propagation, incomplete curation, and missing experimental validation, the overannotation of paralogs further complicates functional annotation in microbial data (88).

In this work, we addressed these challenges by establishing a protein embedding-based targeted function discovery (PE-TFD) framework that explicitly incorporated both positive and negative training data to define a functional search space. Rather than assigning definitive annotations, PE-TFD is designed to generate curated candidate sets enriched for proteins likely to share a target function, even in the absence of detectable sequence or structural similarity.

In a case study with hydrogenases, PE-TFD identified a substantial number of candidates not recovered by sequence-, profile-, or structure-based methods. For NiFe hydrogenases, this included 37% of the proteins lacking any prior functional annotation. While these predictions cannot replace experimental validation, they provide a focused starting point for downstream analyses and the establishment of computational and experimental feedback loops (85). Importantly, PE-TFD is not limited to hydrogenases but can be adapted to other protein families by curating the training data using domain-specific expertise.

Overall, PE-TFD demonstrated that the combination of protein embeddings with discriminative ML/DL strategies offers a shift in perspective from annotation transfer toward candidate prioritization, thereby providing a practical approach to navigate the vastness of functional dark matter.

## Supplementary Data

Supplementary Data are available.

## Data Availability

Scripts available at https://zenodo.org/uploads/18095475?token=eyJhbGciOiJIUzUxMiJ9.eyJpZCI6IjBjNjYzNzQ3LTAwNTMtNGYzYS05YjIwLTNlYmZkYzQ0YzNkMyIsImRhdGEiOnt9LCJyYW5kb20iOiI0YzliYTFlMTJlMzI0N2JiM2IwNTZjMTk5ODk5NDVmYyJ9.tiy2xtjC2Q1-EiyL9HTQph6u6EWeiThN1O1GmgnK0k2sOAxUbQ4qLlPsEANOpE5KB5N81keaBl1FCYYKAnhH7Q

## Acknowledgements

The authors thank Dr. Florian Lenk for generating the protein embeddings, Mathias Didio for developing and testing initial feed-forward neural network (FNN) models and Dr. John Vollmers for helpful discussions.

## Authors contributions

S.W. and A.K.K. wrote the manuscript, S.W. did the data curation and designed and performed the computational analyses and A.K.K. conceived the study.

