## SupplementalFiles for "Deciphering functional dark matter: Machine and deep learning-based processing of protein embeddings enables targeted function discoveries"

**Table S1.** Pre-folded protein structures from AlphaFoldDB corresponding to HydDB-listed proteins.

| | | | |
|---|---|---|---|
| AF-A0A0E3M8W6 | AF-D5VR51 | AF-A0LQC1 | AF-I3TEX8 |
| AF-A0A0F7FAG2 | AF-D5XCK6 | AF-A1AUU6 | AF-I3VT58 |
| AF-A0A0T6LSA8 | AF-J4XFA3 | AF-A1HLY9 | AF-I3XST7 |
| AF-A0A0U1HTU5 | AF-J7ITC3 | AF-A3CSE3 | AF-I3ZUH0 |
| AF-A0A0U1KUH9 | AF-J7QQM9 | AF-A4CGP4 | AF-I7KZA0 |
| AF-A0A0W0SAE2 | AF-K0YJ11 | AF-A4EBM0 | AF-I7LGK1 |
| AF-A0A0W1DJ43 | AF-K4LIP8 | AF-A5GAS4 | AF-J0SAL0 |
| AF-A0A1B4LJJ8 | AF-K4RFV2 | AF-A6DBV4 | AF-J1L1M5 |
| AF-A0A1C0W1X0 | AF-M2ZAC9 | AF-A6Q3L5 | AF-J1L3A3 |
| AF-A0A1D2WF45 | AF-M5TCZ0 | AF-A7I9X6 | AF-A0A7U3YN00 |
| AF-A0A1D2WI80 | AF-M9SHJ3 | AF-A8IMR1 | AF-A8UUB0 |
| AF-A0A1I3NFP2 | AF-N0BGR8 | AF-A8UU10 | AF-D0IAG9 |
| AF-A0A1X1MVN9 | AF-O26498 | AF-B1I3Q5 | AF-F2L646 |
| AF-A0A2S9VD12 | AF-O66988 | AF-B1IMA8 | AF-F2LWC9 |
| AF-A0A3R5YBT8 | AF-Q0W2X7 | AF-B1L3Q0 | AF-F2NCW5 |
| AF-A0A5B9TR73 | AF-Q1MR82 | AF-D6SSI4 | AF-G2FTE3 |
| AF-A0A6B8M4T6 | AF-Q2IN69 | AF-D6Z0J7 | AF-G9XP26 |
| AF-A0A6G6IW48 | AF-Q2LYD7 | AF-E3D146 | AF-H0PT27 |
| AF-A0A6N8JQ21 | AF-Q3Z8U2 | AF-D6SSB2 | AF-D6Z3I9 |
| AF-A0A7T4KNN0 | AF-B3E9J3 | AF-F6BBF0 | AF-E0UTJ2 |
| AF-Q3Z856 | AF-B3QKV3 | AF-F6BE61 | AF-E1QGS6 |
| AF-Q6AL34 | AF-B4UIS3 | AF-F6D3D6 | AF-E1RHQ5 |
| AF-Q8PUL0 | AF-B5EBV8 | AF-F6DVL0 | AF-E3GYP4 |
| AF-Q8TRN3 | AF-B5Y915 | AF-F7XLB9 | AF-E4S640 |
| AF-Q21QY4 | AF-B5YKR2 | AF-F8AF44 | AF-E4U2U9 |
| AF-Q21R14 | AF-B6BP18 | AF-F8AJ81 | AF-E6WY97 |
| AF-Q21RP9 | AF-B9L8N1 | AF-F8E9Q5 | AF-E8MXZ8 |
| AF-Q30QL7 | AF-C0GED6 | AF-F8KT56 | AF-E8T2I8 |
| AF-Q30ZG1 | AF-C0QP83 | AF-F8WSV8 | AF-E8T5B2 |
| AF-Q65PZ2 | AF-C0QP88 | AF-G0HL37 | AF-M1P8A7 |
| AF-Q729Q8 | AF-C0QPV5 | AF-G0HM74 | AF-E8T5B8 |
| AF-R4G257 | AF-C1DUE7 | AF-G0JU97 | AF-F0LMZ5 |
| AF-R9SJG8 | AF-C1DUT5 | AF-G1WIL8 | AF-F0S2N6 |
| AF-S5ZDV0 | AF-C5CEV3 | AF-G1WL42 | AF-F0S3G9 |
| AF-S5ZWS7 | AF-C7MNR6 | AF-G2E383 | AF-H1KZP7 |
| AF-D3DGG7 | AF-C7NU24 | AF-H1XT82 | AF-K9HK98 |
| AF-D3DGK9 | AF-C9LQZ6 | AF-H2C1C5 | AF-M1E641 |
| AF-D3DI11 | AF-C9R996 | AF-H2FWX0 | AF-M1WJX5 |
| AF-D3E0C7 | AF-C9RFJ5 | AF-H3ZK60 | AF-S7T3C6 |
| AF-D3F8J9 | AF-A0A7U3YQ09 | AF-H5WXX8 | AF-S7UEQ0 |
| AF-D3FUC5 | AF-A0A7U4DPF4 | AF-H7C6M7 | AF-U5NA68 |
| AF-D3RMN4 | AF-A0A7U8C660 | AF-H8I6J0 | AF-V6L5F4 |
| AF-D3RWZ6 | AF-A0A101QDC6 | AF-H8I7J4 | AF-W0DCT0 |
| AF-D3SMR5 | AF-A0A174AHV1 | AF-H8I9E5 | AF-W0DGE7 |
| AF-D5CSB4 | AF-A0A328ETC9 | AF-H8I539 | |
| AF-D5U199 | AF-A0A376D7X9 | AF-I3CK80 | |
| AF-D5V582 | AF-A0A806X6S7 | AF-I3REP4 | |

**Table S2.** Cut-off values (best domain) for public HMM-based profiles.

| | | |
|---|---|---|
| **[FeFe] hydrogenase** | | |
| ***Excellent candidate*** | | |
| | TIGR02512.1 | > 400 |
| AND/OR | TIGR04105.1 | > 400 |
| AND/OR | PF02906.18 | > 350 |
| NOT | PTHR11615.SF124 | > 350 |
| NOT | PTHR11615.SF258 | > 350 |
| ***Good candidate*** | | |
| | TIGR02512.1 | > 200 |
| AND/OR | TIGR04105.1 | > 200 |
| AND/OR | PF02906.18 | > 200 |
| NOT | PTHR11615.SF124 | > 350 |
| NOT | PTHR11615.SF258 | > 350 |
| ***Soft candidate*** | | |
| | TIGR02512.1 | > 70 |
| AND/OR | TIGR04105.1 | > 70 |
| AND/OR | PF02906.18 | > 70 |
| NOT | PTHR11615.SF124 | > 350 |
| NOT | PTHR11615.SF258 | > 350 |
| **[NiFe] hydrogenase** | | |
| ***Excellent candidate*** | | |
| | NF033181.1 | > 250 |
| AND/OR | NF040828.1 | > 250 |
| AND/OR | NF040832.1, | > 250 |
| AND/OR | NF041785. | > 250 |
| AND/OR | TIGR03295.1 | > 100 |
| AND/OR | PTHR42958 | > 350 |
| AND/OR | or PF00374.23 | > 250 |
| NOT | PF00346.23 | > 250 |
| NOT | PTHR11993 | > 250 |
| NOT | TIGR01962.1 | > 250 |
| ***Good candidate*** | | |
| | NF033181.1 | > 150 |
| AND/OR | NF040828.1 | > 150 |
| AND/OR | NF040832.1 | > 150 |
| AND/OR | NF041785.2 | > 150 |
| AND/OR | TIGR03295.1 | > 85 |
| AND/OR | PTHR42958 | > 250 |
| AND/OR | PF00374.23 | > 100 |
| NOT | PF00346.23 | > 250 |
| NOT | PTHR11993 | > 250 |
| NOT | TIGR01962.1 | >250 |
| ***Soft candidate*** | | |
| | NF033181.1 | > 50 |
| AND/OR | NF040828.1 | > 50 |
| AND/OR | NF040832.1 | > 50 |
| AND/OR | NF041785.2 | > 50 |
| AND/OR | TIGR03295.1 | > 50 |
| AND/OR | PTHR42958 | > 100 |
| AND/OR | PF00374.23 | > 50 |
| NOT | PF00346.23 | > 75 |
| NOT | PTHR11993 | > 100 |
| NOT | TIGR01962.1 | > 100 |

**Table S3.** Positive and negative training data sets for [NiFe] and [FeFe] hydrogenases.

{Excel table}

**Table S4.** Shannon indices.

| Training set | Protein identity in cluster | Number of clusters | Shannon index H' | Maximal Shannon index H'max | Evenness |
|---|---|---|---|---|---|
| FeFe | 0.3 | 7 | 1.1634 | 1.9459 | 0.5979 |
| FeFe | 0.5 | 69 | 3.1316 | 4.2341 | 0.7396 |
| FeFe | 0.7 | 391 | 5.6016 | 5.9687 | 0.9385 |
| FeFe | 0.9 | 683 | 6.4144 | 6.5265 | 0.9828 |
| FeFe | 0.95 | 748 | 6.5604 | 6.6174 | 0.9914 |
| NiFe | 0.3 | 4 | 0.4494 | 1.3863 | 0.3242 |
| NiFe | 0.5 | 42 | 2.5349 | 3.7377 | 0.6782 |
| NiFe | 0.7 | 222 | 4.6525 | 5.4027 | 0.8611 |
| NiFe | 0.9 | 679 | 6.3566 | 6.5206 | 0.9748 |
| NiFe | 0.95 | 794 | 6.5929 | 6.6771 | 0.9874 |

**Table S5.** Hyperparameters for [NiFe] hydrogenase training [FeFe] hydrogenase training.

| | |
|---|---|
| **Gaussian Naive Bayes (NB)** | |
| [NiFe] hydrogenase | var_smoothing: 0.0029449109184381457 |
| [FeFe] hydrogenase | var_smoothing: 0.0006428708289462023 |
| **Logistic Regression (LR)** | |
| [NiFe] hydrogenase | C: 2.93478126621975, solver: liblinear |
| [FeFe] hydrogenase | C: 13.244885037820286, solver: liblinear |
| **Support Vector Machine (SVM)** | |
| [NiFe] hydrogenase | C: 0.008973293313175697, kernel: rbf |
| [FeFe] hydrogenase | C: 9.096622199069632, kernel: rbf |
| **Random Forest (RF)** | |
| [NiFe] hydrogenase | n_estimators: 77, max_dept: 4, min_samples_split: 14 |
| [FeFe] hydrogenase | n_estimators: 30, max_depth: 9, min_samples_spli: 11 |
| **K-nearest neighbor (KNN)** | |
| [NiFe] hydrogenase | neighbors: 12 |
| [FeFe] hydrogenase | neighbors: 2 |
| **Feed-forward neural network (FNN)** | |
| [NiFe] hydrogenase | num_hidden_layers: 2, hidden_size_0: 90, hidden_size_1: 112, dropout_rate: 0.16950096904139747, learning_rate: 0.0017258039402130614, optimizer: Adam, batch_size: 42, epochs: 11 |
| [FeFe] hydrogenase | num_hidden_layers: 3, hidden_size_0: 126, hidden_size_1: 97, hidden_size_2: 118, dropout_rate: 0.2569864949338112, learning_rate: 0.0006595636906350461, weight_decay: 6.853729752935781e-06, optimizer: Adam, batch_size: 40, epochs: 13 |

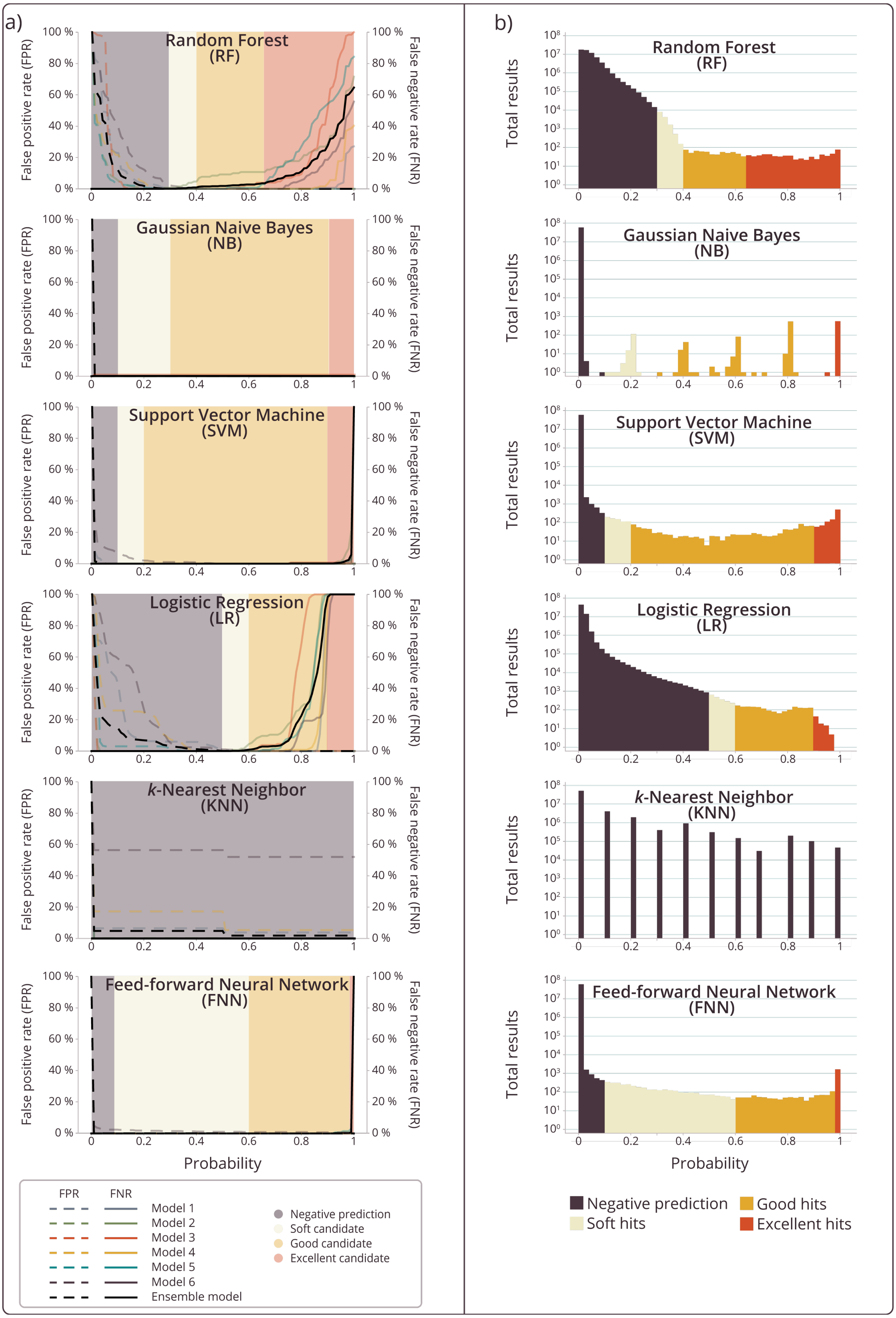
a)
Random Forest (RF)
Gaussian Naive Bayes (NB)
Support Vector Machine (SVM)
Logistic Regression (LR)
k-Nearest Neighbor (KNN)
Feed-forward Neural Network (FNN)
False positive rate (FPR)
False negative rate (FNR)
Probability
FPR
FNR
Model 1
Model 2
Model 3
Model 4
Model 5
Model 6
Ensemble model
Negative prediction
Soft candidate
Good candidate
Excellent candidate
b)
Total results
Probability
Negative prediction
Soft hits
Good hits
Excellent hits

**Figure S1. Performance of each classifier during testing and prediction - [FeFe] hydrogenases. a)** False positive rates (FPR) and false negative rates (FNR) across probability thresholds. Each panel shows the FPR (dashed lines, left y axis) and FNR (solid lines, right y axis) for models 1 to 6 across probability thresholds for one of the applied classifiers. In addition, the rates of the ensemble model (models 1,2,3,4,5) are given. The curves inform the selection of probability thresholds that balance sensitivity and specificity for identifying novel [FeFe] hydrogenase candidates. **b)** Number of total prediction results across probability thresholds. Distribution of predicted probabilities for all proteins in UniRef50 generated by each PE-TFD classifier. The background shading for a) and b) indicates the probability ranges used to define prediction categories: negative prediction (red), *soft* candidates (light orange), *good* candidates (yellow), and *excellent* candidates (green). Given the odd result structure, KNN results were not used further.

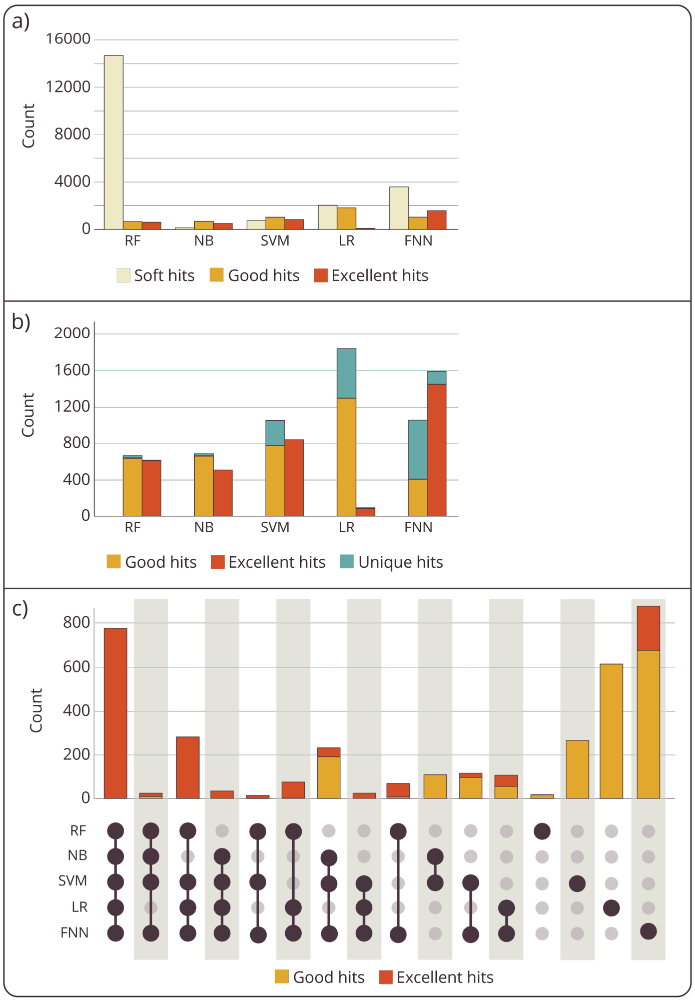


**Figure S2 PE-TFD classifier performance on [FeFe] hydrogenase-aimed predictions. a)** Numbers of *soft*, *good*, and *excellent* candidates predicted by each classifier. **b)** Proportion of candidates detected exclusively by each classifier. **c)** Intersection analyses of good and excellent candidates and the six applied ML/DL classifiers. Combinations with fewer than 10 proteins were omitted for clarity.

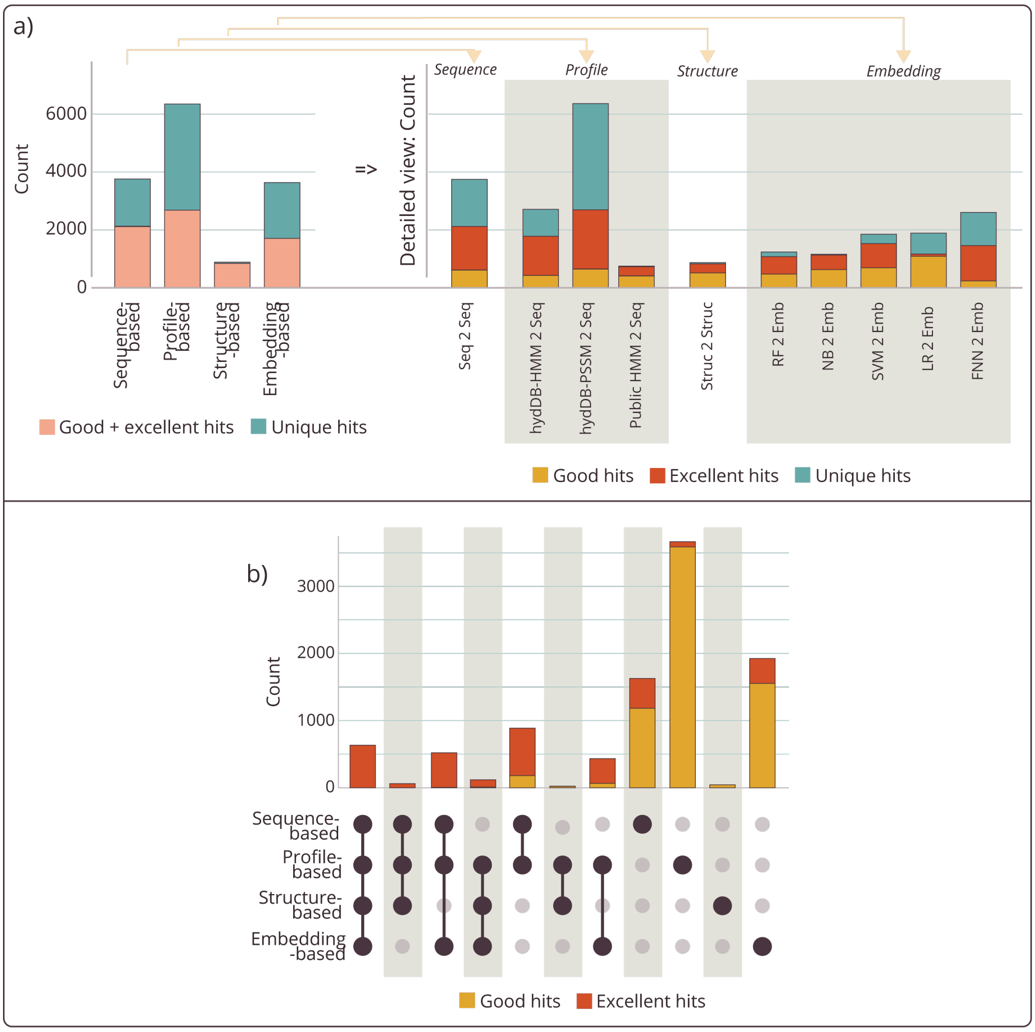


**Figure S3. Benchmarking PE-TFD results against sequence-, profile-, and structure-based approaches – [FeFe] hydrogenases. a)** Proportion good and excellent candidates produced by different method categories, including sequence-based (*Seq2Seq*), profile-based (*HydDB-HMM2Seq*, *HydDB PSSM2Seq*, *Public HMM2Seq*), structure-based (*Struc2Struc*), and protein embedding-based (RF, NB, SVM, LR, FNN) approaches. KNN predictions were not included in the analysis. **b)** Intersection analyses of good and excellent candidates within PE-TFD and benchmarking results.

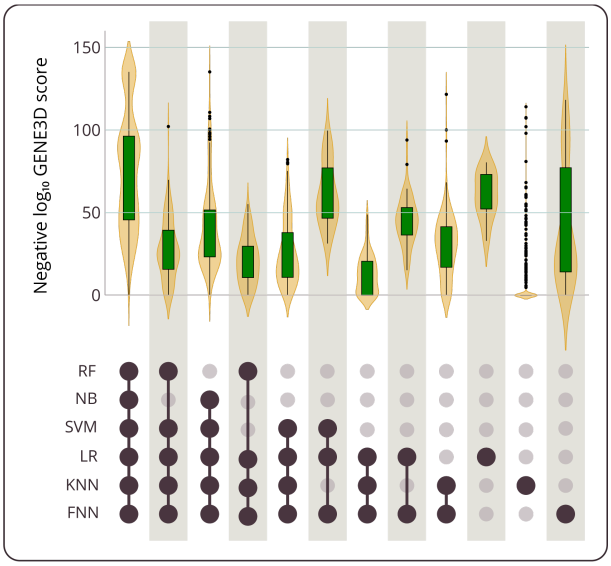


**Figure S4 PE-TFD classifier performance on [NiFe] hydrogenase-aimed predictions.** Combinations with fewer than 5 proteins were omitted for clarity. Confidence scores of hydrogenase-associated structural domains of the GENE3D database in PE-TFD candidates and benchmarks.

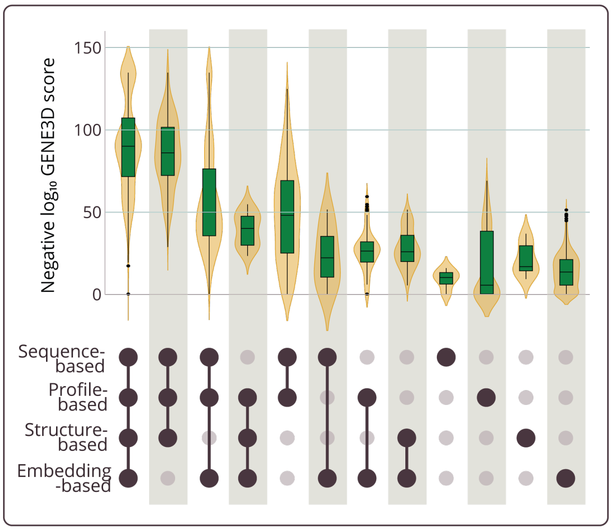


**Figure S5 Intersections and structural breakdown of PE-TFD and benchmarking results – [NiFe] hydrogenase.** Combinations with fewer than 5 proteins were omitted for clarity. Confidence scores of hydrogenase-associated structural domains of the GENE3D database in PE-TFD candidates and benchmarks.

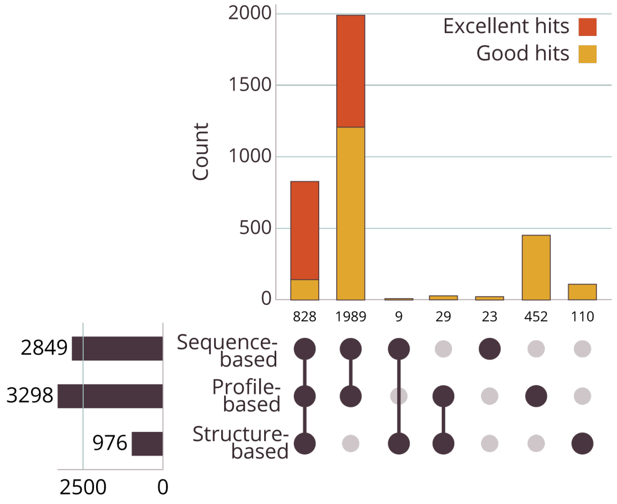


**Figure S6 Intersection analysis of PE-TFD and 3D structure results – [NiFe] hydrogenase.** The results are depicted without the PE-TFD results.

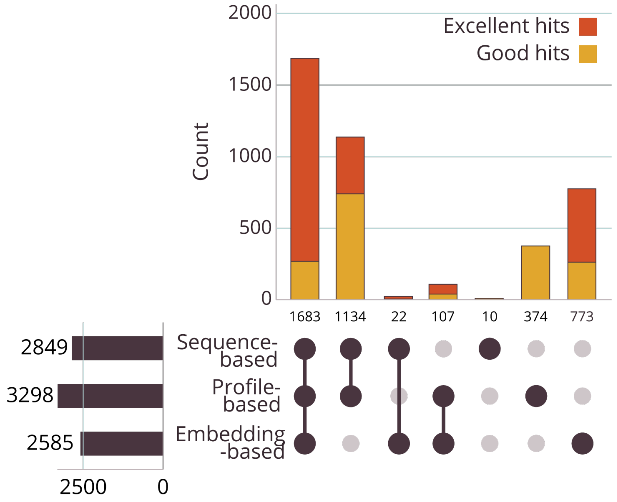


**Figure S7 Intersection analysis of PE-TFD and goldstandard results – [NiFe] hydrogenase.** The results are depicted without the results for the structural analysis.
